# *In silico* identification and deorphanisation of an allatostatin C GPCR system in the cephalopod *Octopus vulgaris* reveals two receptors with distinct potency

**DOI:** 10.64898/2026.02.18.706622

**Authors:** Eleonora Maria Pieroni, James Dillon, Vincent O’Connor, Lindy Holden-Dye, Pamela Imperadore, Graziano Fiorito, Luis Alfonso Yañez-Guerra

**Affiliations:** School of Biological Sciences, University of Southampton, Southampton SO17 1BJ, UK; Association for Cephalopod Research ‘CephRes’ ETS, Napoli, Italy; Department of Biology and Evolution of Marine Organisms, Stazione Zoologica Anton Dohrn, Napoli, Italy

**Keywords:** neuropeptides, calcium assay, evolution, deorphanisation

## Abstract

Neuropeptide signalling is transversally important in all living animals as it constitutes the basis of cellular communication. The investigation of the functional roles of peptide signalling represents an important route to understanding evolution of specific physiological traits and behaviours in metazoans.

Allatostatins and their cognate receptors are classically defined as invertebrate neuropeptide hormones. Among these, allatostatin C was firstly associated with insect development. However, accumulating evidence recognises the presence of allatostatin C as a conserved signalling molecule across all invertebrate lineages, with reported functions spanning from regulation of feeding and digestion to immune responses and modulation of core nociception.

Here we combined *in silico* and experimental approaches to reveal the interacting molecular determinants of the allatostatin C signalling in the cephalopod *Octopus vulgaris,* a scientifically and culturally interesting invertebrate for its centralised nervous system, capable of top-down modulation of complex behaviours. This resolved a single prepropeptide encompassing allatostatin C peptide (OvAstC), whose conserved mature form (AVITACYFQAVSCY) was shown to differentially activate two identified cognate receptors (OvAstCR1 and OvAstCR2) when heterologously expressed in the recombinant system HEK293G5A.

PCR analysis carried out in *O. vulgaris* tissues, showed a broad distribution of OvAstC and OvAstCRs. This wide expression across nervous, immune and digestive tissues is consistent with a pleiotropic role of this peptidergic system. Together, the opioid/somatostatin-related phylogenetic placement of OvAstCRs and the broad expression of OvAstC components in nervous and sensory tissues nominate this pathway as a candidate for neuromodulatory control of sensory processing, including nociception, with potential welfare relevance in cephalopods.

**Significance statement:** Cephalopods represent an evolutionarily distinctive molluscan lineage that evolved a centralised nervous system capable of displaying advanced learning and behavioural complexity compared with other invertebrates. These features, speculated to allow elaboration of pain-like states, granted cephalopods inclusion as the only invertebrate taxon requiring protection under European legislations when used in research. Investigation of the neuropeptide signalling in cephalopods is currently understudied despite its crucial role in regulating broad physiological functions in organisms. This study identified for the first time a single allatostatin C peptide and two cognate receptors in *Octopus vulgaris*. Our characterisation of a putative endogenous allatostatin C system in octopus, the accumulating evidence of its central role in invertebrate antinociception and its evolutionary relationship with the vertebrate-exclusive analgesic opioid family, represent a critical starting point for a more in-depth analysis of the physiological role of allatostatin C in this subclass of molluscs, with important welfare implications.

## Introduction

Neuropeptide signalling plays a pivotal role in modulating animal physiology and function, including the integration and modulation of stimuli coming from the exteroceptive and interoceptive environmental cues. This broad biological significance is reinforced by evidence of peptidergic signalling across all animals, including “neuron-less” organisms such as sponges and placozoans (Hauser, Koch and Grimmelikhuijzen, 2022; Yañez-Guerra, Thiel and Jékely, 2022).

Classically, identifying peptide homologues across animal phyla was rather difficult and challenged by the short and divergent nature of these molecules (Veenstra, 2010). However, in the last decade, clustering-based approaches in conjunction with large-scale de-orphanisation, have contributed to the dissection of numerous neuropeptides and their cognate receptors across different phyla, including those belonging to key phylogenetic lineages that are informative for understanding the evolution of peptidergic signalling in animals such as annelids or cnidarians (Bauknecht and Jékely, 2015; Thiel *et al.*, 2024).

Allatostatin C is a neuropeptide originally identified as an inhibitor of the juvenile hormone biosynthesis in the corpora allata of the tobacco hornworm *Manduca sexta,* leading to developmental changes such as metamorphosis and sexual maturation (Kramer *et al.*, 1991). Subsequent studies have revealed roles in insect feeding and digestion, modulation of immune responses and downstream mitigation of nociception (Ma *et al.*, 2009; Duan Sahbaz *et al.*, 2017; Bachtel *et al.*, 2018; Zhang *et al.*, 2022; Liu *et al.*, 2023). These pivotal functions have heightened pharmacological interest in the potential of allatostatin C signalling system as a suitable pest control target (Işbilir *et al.*, 2020; Shahraki *et al.*, 2021).

Allatostatin C was found to be conserved in other invertebrates where it exerts distinct tissue-selective functional roles (Audsley and Weaver, 2007; Stay and Tobe, 2007; Ma *et al.*, 2009; Bachtel *et al.*, 2018; Kubrak *et al.*, 2022; Zhang *et al.*, 2022). Characterisation of this system in molluscs largely relied on *in silico* analyses and tissue expression studies, such as in the gastropod *Lottia gigantea*, (Veenstra, 2010) and the bivalve *Mizuhopecten yessoensis* (Zhang *et al.*, 2024), but these studies did not investigate the functional activation of the identified receptors by their candidate ligands.

The only allatostatin C receptor de-orphanisation study conducted on a mollusc was carried out in *Aplysia californica*, where an interaction between the identified receptor and allatostatin C peptide was experimentally demonstrated (Jiang *et al.*, 2022).

Despite their complex brain anatomy and associated physiology, which imply extensive peptidergic modulation, functional investigation of neuropeptide signalling in cephalopods remains limited.

Previous studies have reported functional characterisation of the Gonadotropin-Releasing Hormone (GnRH), tachykinin and octopressin/cephalotocin peptides in the common octopus *O. vulgaris* through heterologous expression in the recombinant *Xenopus laevis* system (Kanda *et al.*, 2005; Kanda *et al.*, 2006; Kanda *et al.*, 2007), but no de-orphanisation work was carried out on the allatostatins.

To date, only one study has investigated the neuropeptidome landscape of the cuttlefish *Sepia officinalis* using RNA sequencing, *in silico* analysis and tissue mapping through mass spectrometry, but failed to identify allatostatin C (Zatylny-Gaudin *et al.*, 2016).

Here we report de-orphanisation of the putative allatostatin C system in the cephalopod mollusc *O. vulgaris*.

Using *in silico* approaches and experimental validation, we detected two *bona fide* allatostatin C receptors, OvAstCR1 and OvAstCR2, belonging to a class A GPCR cluster that includes opioid (OPR), somatostatin (SST) and melanocortin (MCR) receptors. A single allatostatin C (OvAstC) peptide (AVITACYFQAVSCY) was identified from *O. vulgaris* genome and demonstrated to act on both receptors, although with different potency. Importantly, we identified a broad tissue distribution of all the molecular determinants of octopus allatostatin C signalling which suggests a wide range of physiological functions (e.g., immune-, digestive and sensory related) in a way that overlaps with the evolutionary related deuterostome- and vertebrate-specific somatostatin and opioid receptor families respectively (Krantic, 2000; Olias *et al.*, 2004; Sternini *et al.*, 2004). From an evolutionary perspective, these findings position cephalopods as a key comparative lineage for understanding the diversification and functional specialisation of ancient peptidergic signalling systems in bilaterians. This would provoke the investigation of physiological functions of these peptide signalling clusters that are overlooked or difficult to study but which are leveraged by characterising potential molecular determinants, as exemplified by pathways implicated in nociception. Given the significance of this biological domain in cephalopods, our identification of the putative endogenous representatives of the AstC system, provides pharmacological targets that could be further characterised for their role in pain-related physiology, supporting attention to studying peptidergic signalling in cephalopods as part of a broader evolutionary and welfare context.

## Results

### Characterisation of *Octopus vulgaris* allatostatin-C receptors reveals conserved GPCR features and evolutionary links to somatostatin and opioid receptors

In our previous *in silico* analysis of candidate *O. vulgaris* nociceptive related genes, we identified two potential allatostatin C receptors: c220802_g3_i1 (<u>PV164564.1</u>) and c31154_g2_i1 (<u>PV164563.1</u>) here after referred as *OvAstCR1* and *OvAstCR2*, respectively (Pieroni *et al.*, 2026). When a more complete *O. vulgaris* genome became available (Destanović *et al.*, 2023), we further analysed the transcript sequences to confirm the presence of these receptors and potentially identify new candidates.

By checking the automated protein predictions associated with the new *O. vulgaris* genome (Destanović *et al.*, 2023), we found *OvAstCR1* completely matched with OctVul6B022141P1 encoding for a 410 amino acidic sequence, whilst *OvAstCR2* corresponded to OctVul6B016349P which was originally incomplete (195 amino acids) due to a mispredicted start codon when compared to the c31154_g2_i1 (443 amino acids). The secondary structure of the amino acids predicted for the two receptors genes showed the classical G-protein coupled receptors (GPCRs) characteristics. This included an extracellular N-terminal, 7 transmembrane elements (TMs) and an intracellular C- terminal (Figure 1). The two receptors showed an overall 47.8% identity (64.5% similarity) between each other, and a 56.5% identity (71.7% similarity) when considering only the region spanned by the characteristic seven TMs.

**Figure 1.**
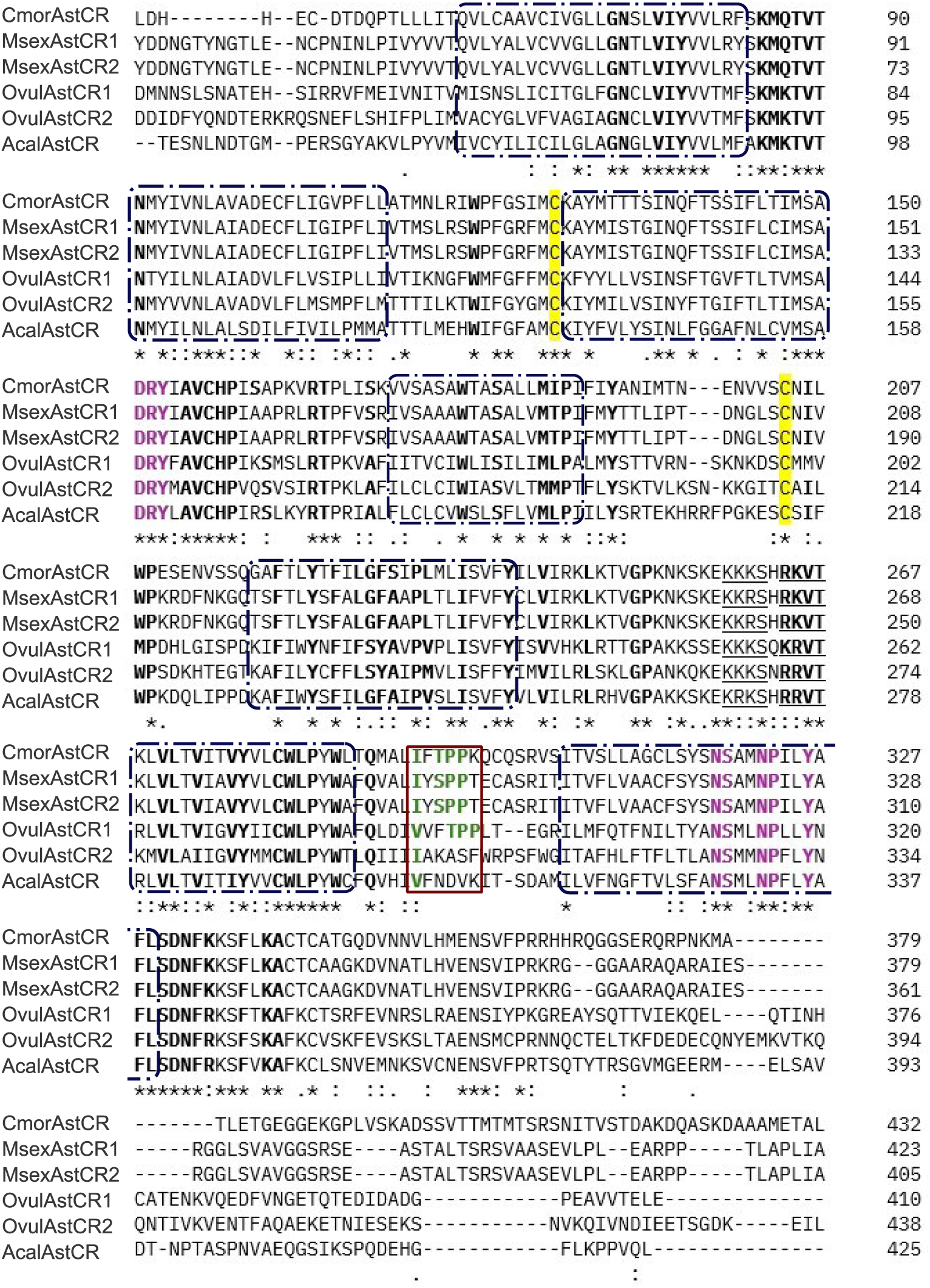
Sequence alignment highlights class A GPCR conserved residues in *O. vulgaris* putative allatostatin C receptors. Alignment of OvAstCR1 and OvAstCR2 with AstCRs of indicated invertebrate species. The alignment revealed key GPCR A signatures: the seven transmembrane elements (dash-pattern frames), conserved functional residues (in bold black), the residues for functional activation of the receptor (in bold magenta) and the cysteine residues for the disulphide bond formation (highlighted in yellow). Predicted cAMP and cGMP-dependent phosphorylation sites (KKKS, RKVT) found in somatostatin receptors are underlined. The proposed key residues implicated in the ligand binding region in the third extracellular loop (Duan Sahbaz *et al.,* 2017) are shown in green and the conserved motif IxPP is framed in red. Interestingly, a similar but slightly different VxxPP was found in the OvAstCR1 but no conserved residues are found in OvAstCR2. Cmor: *Carausius morosus* (KX255655.1); Msex1: *Manduca sexta* AstCR1 (A0A921ZMA1); Msex2: *Manduca sexta* AstCR2 (XP_030033910_1); Acal: *Aplysia californica* (XP_005095139_1).

Signature residues such as the DRY, NPILY and WLP motifs as well as the cysteines involved in a disulfide bond formation in the extracellular loop (ECL) 2 allowed the classification of these predicted receptors as members of the Rhodopsin family (class A GPCR) representatives (Figure 1). Interestingly, the putative cAMP- and cGMP-dependent phosphorylation motifs KKKS and xxVT, as well as conserved residues in the TM7 that are typically found in the orthologous deuterostome-specific somatostatin receptors were also identified (Figure 1), thus supporting the hypothesis these receptors were allatostatin C receptors (Duan Sahbaz *et al.*, 2017).

Next, we carried out a cluster and phylogenetic analysis using reference sequences from well-characterised and curated invertebrate allatostatin C receptors and vertebrate somatostatin (SSRs) and opioid receptors (OPRs), as these classes of GPCRs were previously shown to be related (Mirabeau and Joly, 2013). We performed an initial BLAST search against the complete proteome of 21 representative species from different phyla (Supplementary File S1) and visualised the sequences relationship through a cluster analysis using CLANS (Zimmermann *et al.*, 2018). The CLANS map showed numerous representatives of the Rhodopsin GPCR class, such as monoamine receptors, opsins, and neuropeptide F and FF receptors (Figure 2). The two *O. vulgaris* sequences of interest OvAstCR1 and OvAstCR2 were found to be the only *O. vulgaris* receptors within a cluster with strong connections with AstCRs, SSRs and OPRs (Figure 2). Additionally, these sequences showed connections within the cluster with melanocortin receptors (MCRs), galanin receptors (GALRs), Kisspeptin 1 receptors (Kiss1Rs) and the invertebrate allatostatin A receptors (AstARs). All the sequences connected within the same cluster (197 sequences), were used to build a phylogeny tree and resolve their relationship more in detail.

**Figure 2.**
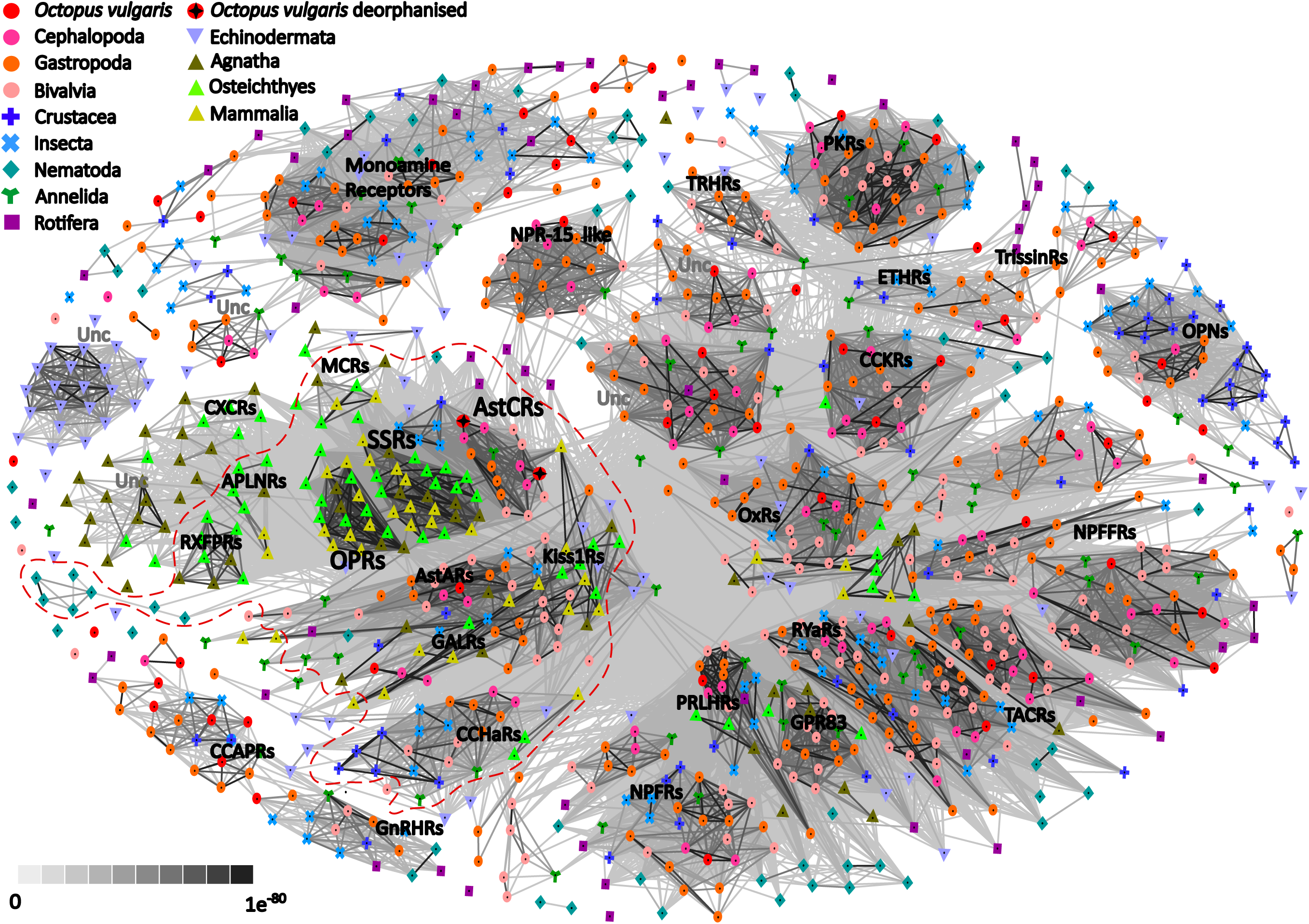
OvAstCR1 and OvAstCR2 are allatostatin C receptors with evolutionary relationship with somatostatin and opioid receptors. Following a BLAST search for AstCRs, SSRs, OPRs, and related class A GPCRs across different representative species, a CLANS analysis has been carried out to reveal the relationship between each other. Each dot represents a species-specific receptor sequence (see legend in top left corner). The gradient of the connection represents the degree of similarities between sequences (indicated in the bottom left corner). The analysis shows the identified *O. vulgaris* receptors OvAstCRs are indeed Allatostatin C receptors and share the same cluster with somatostatin and opioid receptors (dashed red frame). APLNR: apelin receptor, AstAR: allatostatin A receptor, AstCR: allatostatin C receptor, CCAPR: crustacean cardioactive peptide receptor CXCR: CCHaR: CChamide receptor, CCKR: cholecystokinin receptor, CXC chemochine receptor, ETHR: ecdysis triggering hormone receptor, GALR: galanin receptor, GnRHR: gonadotropin releasing hormone receptor, GPR83: G-protein coupled receptor 83, Kiss1R: Kisspeptin 1 receptor, MCR: melanocortin receptor, NPFFR: neuropeptide FF receptor, NPFR: neuropeptide F receptor, NPR-15: neuropeptide 15 like receptor, OPN: opsin receptor, OPR: opioid receptor, OxR: orexin receptor, PKR: pyrokinin receptor, PRLHR: Prolactin-Releasing Hormone Receptor, RYaR: RYamide receptor, RXFPR: relaxin family peptide receptor, SSR: somatostatin receptor, TACR: tachykinin receptor, TRHR: thyrotropin-releasing hormone receptor, TrissinR: trissin receptor, Unc: uncharacterised

The phylogenetic analysis showed that AstCRs are an evolutionary related but distinct class of invertebrate receptors from the vertebrate-exclusive OPRs. Additionally, as previously shown, the phylogeny is consistent with a duplication event that led to the AstCR and the deuterostome-specific SSRs branches (Figure 3). Specifically looking at *O. vulgaris*, the two sequences of interest were identified as part of the allatostatin C branch, confirming their identity as allatostatin C receptors (Figure 3).

**Figure 3.**
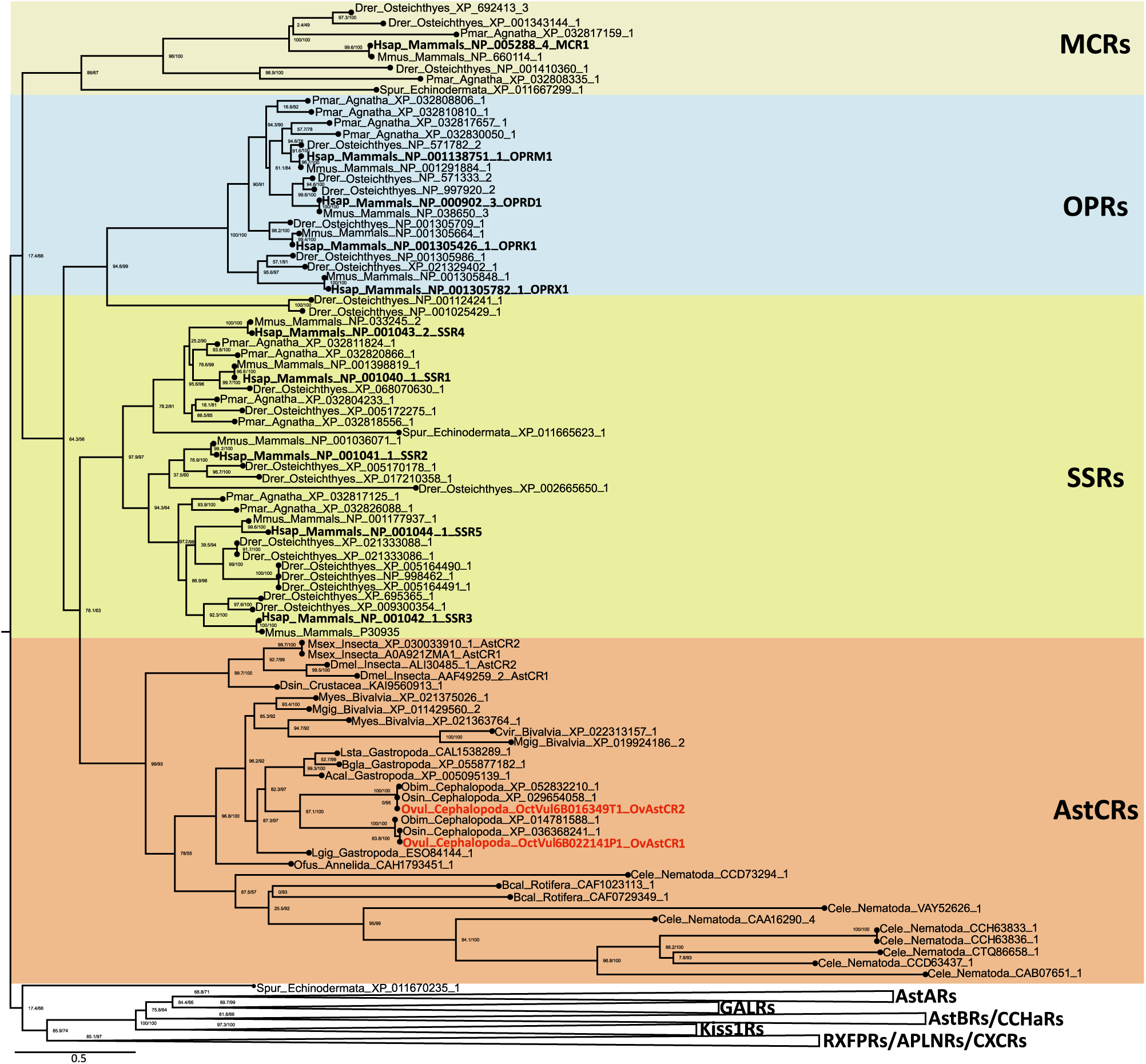
Phylogenetic analysis reveals OvAstCRs are allatostatin C receptors. Allatostatin C receptors (red section) are the closest protostome orthologues of the vertebrate exclusive OPRs (blue section) and paralogues of the deuterostome-specific SSRs (yellow section). The OvAstCRs are highlighted in bold red and are clearly part of the AstCRs branch. The phylogenesis has been carried out using the 197 sequences belonging to the same identified cluster (Figure 2). The tree has been rooted using the outgroup sequences such as GALRs, AstARs, Kiss1Rs. All sequences used are available in Supplementary File S3.

### Two receptors, one peptide: *O. vulgaris* has one putative endogenous AstC peptide

Next, we performed a BLAST search against *O. vulgaris* genome in search of the candidate endogenous activators of the OvAstCRs (Destanović *et al.*, 2023) using AstC precursor sequences from different protostomes (Uniprot Taxon id: 33317). The search produced a single hit (OctVul6B022881P1) with an identified 22 amino acid signal peptide sequence and a typical dibasic KR cleavage site with a single embedded 14 amino acid peptide sequence (Figure 4A). When aligned with other invertebrate AstC precursor sequences, highly conserved residues within the predicted mature peptide sequence such as the two cysteines, known to classically cyclise AstC peptides via the formation of a disulfide bond, were revealed (Figure 4B). The amino acid residues surrounding the cysteines shared the strongest conservation. The only exception was the third residue after the first cysteine, in which a conserved asparagine (N) was found in all protostome phyla expect for cephalopods. In this group, this position is occupied by a glutamine (Q), with 100% identity observed in the mature peptide sequence (Figure 4B).

**Figure 4.**
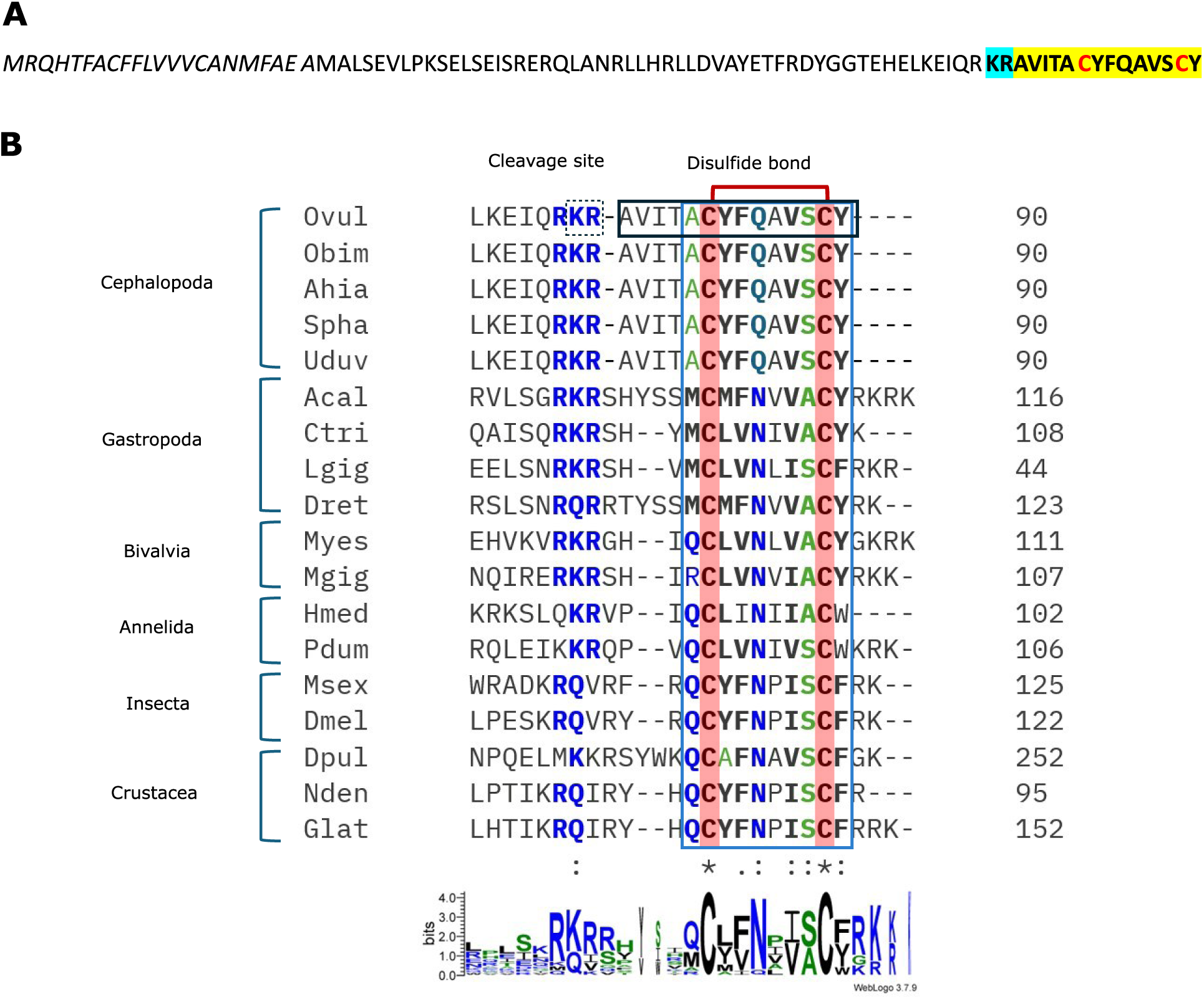
Precursor sequence of *O. vulgaris* allatostatin C peptide and alignment with allatostatin C precursors from different invertebrate phyla. **A** *In silico* analysis predicted the presence of one *O. vulgaris* AstC precursor with a single embedded mature peptide. The signal peptide is underlined. The classical dibasic KR cleavage site is highlighted in cyan, and the putative mature peptide is highlighted in yellow. The two cysteine residues, typically forming a disulfide bridge in the bioactive form of the peptide, are in bold red. **B** The mature AstC shows a high level of conservation across selected representative invertebrate species, particularly in the cysteine residues (here highlighted in red) forming a disulfide bond. Key conserved residues have been highlighted in different colours within the light blue rectangle according to their relative prevalence as depicted by WebLogo 3.7.9 below. The cleavage site KR in the OvAstC (Ovul) sequence is shown by the dashed rectangle, the mature form of the peptide is highlighted by the solid black rectangle. Ovul: *Octopus vulgaris* (OctVul6B022881P1); Obim: *Octopus bimaculoides* (Ocbimv22019504m, from mRNA); Ahia: *Argonauta hians* (GAB1606153.1); Spha: *Sepia pharaonis* (GEIE01009944.1, from TSA); Uduv: *Uroteuthis duvaucelii* (GKZI01071814.1, from TSA) Acal: *Aplysia californica* (XP_005112794.1); Myes: *Mizuhopecten yessoensis* (XP_021356393.1); Mgig: *Magallana gigas* (XP_011412814.1); Ctrit: *Charonia tritonis* (AQS80487.1); Lgig: *Lottia gigantea* (Veenstra, 2010); Dret: *Deroceras reticulatum* (ARS01353.1); Hmed: *Hirudo medicinalis* (FP643552.1 – mRNA sequence); Pdum: *Platynereis dumerilii* (AHB62362.1); Msex: *Manduca sexta* (XP_030040870.1); Dmel: *Drosophila melanogaster* (NP_523542.1); Dpul: *Daphnia pulex* (XP_046455835.1); Nden: *Neocaridina denticulata* (AIY69122.1); Glat: *Gecarcinus lateralis* (A0AA50EVT2 – Uniprot).

### The putative OvAstC system is broadly represented across tissues suggesting different physiological functions

The expression of the authenticated receptors and peptide precursor transcripts was investigated by designing primers spanning the entire length of the transcripts. The amplified products were validated through sequencing.

We additionally aligned the OvAstCRs and OvAstC transcript sequences against the *O. vulgaris* genome to find their correct genomic location (Destanović *et al.*, 2023). *OvAstCR1* (1233 bp) was found on chromosome 23 (Figure 5A) and *OvAstCR2* (1332 bp) was located on chromosome 24 (Figure 5B). Both genes were found to be intronless, whilst the *OvAstC* peptide precursor sequence was found on chromosome 12 and was identified as formed by two exons (102 + 171 bp) and one 24,304 bp intron (Figure 5C).

**Figure 5.**
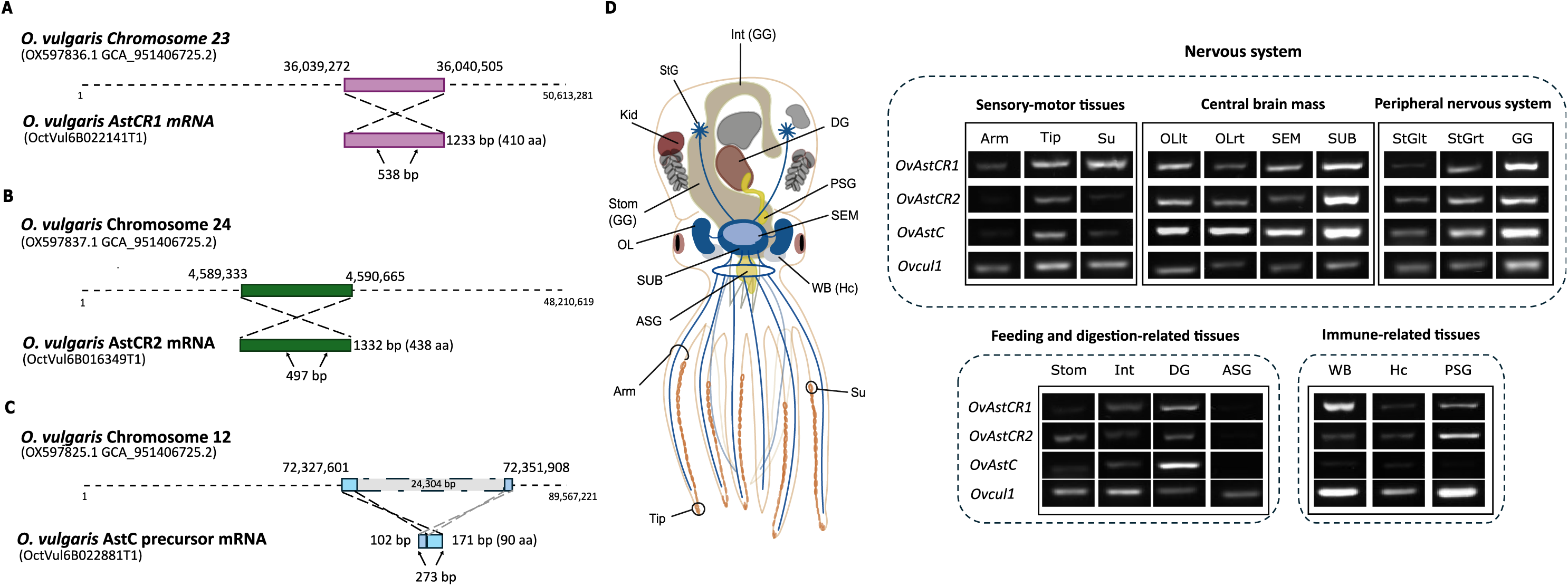
Genomic location and tissue distribution of *OvAstCRs* and *OvAstC* sequences. ***A-C*** Schematic representation of the *O. vulgaris* allatostatin C receptors and related peptide precursor transcripts and their position on the matching chromosome. *OvAstCR1* and *OvAstCR2* are intronless genes found on chromosome 24 and 23. The *OvAstC* precursor transcript is found on chromosome 12 and is composed of two exons and one intron. The arrows underneath the mRNA sequence indicate the region amplified by the designed primers to analyse the tissue distribution. **D** Schematic anatomy of *O. vulgaris* showing the tissues from which RNA was extracted, reverse transcribed and used in the PCR reactions to show the distribution of the genes of interest across key physiological systems. The representative bands shown, refer to PCR performed at least once for each animal tissue (for a total of three animals utilised, Supplementary Table S2) that successfully amplified the control gene *cul1.* This gene was previously shown to be broadly expressed across different *O. vulgaris* tissue types (Imperadore et al., 2023). The amplified bands shown were all of the size expected based on the primer pair selected (see Supplementary Table S3). Arm: arm muscle and embedded axial nerve cord at 50% of its length; Tip: tip of the arm; Su: sucker, OL_lt: left optic lobe; OL_rt: right optic lobe; SEM: supra-oesophageal mass; SUB: sub-oesophageal mass; StG_lt: left stellate ganglion, StG_rt: right stellate ganglion; GG: gastric ganglion; Stom: stomach; Int: intestine; DG: digestive gland; ASG: anterior salivary gland; WB: white body; Hc: haemocytes; PSG: posterior salivary gland.

Following the *in silico* identification and experimental validation of the sequences of the candidate representatives of the *O. vulgaris* AstC system, we proceeded to characterise their expression across a broad set of tissues associated with different physiological functions in octopus (Figure 5D). To this end, we used primers encompassing either the whole (*OvAstC*) or internal (*OvAstCRs*) sequence of the transcript and performed an endpoint PCR using tissues from three individual octopuses.

Our data showed the endogenous peptide precursor is expressed broadly across different tissues, including the central brain mass (i.e., SEM, SUB, OL), the peripheral ganglia (StG, GG) and the sensory-motor tissues (Arm, Tip, sucker). Expression was also found in representative tissues for the feeding and digestion-related (except for the ASG) and immune-related (WB, Hc, PSG) systems (Gestal and Castellanos-Martínez, 2015; Ponte and Modica, 2017; Fernández-Gago, Molist and Rocha, 2019; Figure 5D).

The PCR carried out on *OvAstCR1* and *OvAstCR2* amplified bands of the predicted size (538 bp and 497 bp) which showed an overlapping and broad expression. They were amplified in all the tissues investigated with variable low expression in ASG (Figure 5D).

### OvAstC is the putative endogenous agonist of the OvAstCRs

We next asked whether the candidate genes identified interacted together in a way that would be consistent with a physiological *O. vulgaris* AstC system. To investigate this, we carried out a de-orphanisation of the receptors using the predicted endogenous mature OvAstC peptide. We individually expressed *OvAstCR1* and *OvAstCR2* in the recombinant cell system HEK293G5A, stably expressing the Ca^++^-sensitive aequorin (Baubet *et al.*, 2000). Then the mature peptides were added to record luminescence. This approach is based on exogenous agonist-induced activation of the G_qi_ coupled to the heterologously expressed candidate receptor that leads to an increase in intracellular Ca^++^ levels which subsequently activates aequorin to produce a luminescence signal (Thiel *et al.*, 2024; Yañez-Guerra *et al.*, 2025).

We synthesised the mature peptide (AVITACYFQAVSCY) both in the open (OvAstC_nc) and cyclised structure (with a disulfide bridge between the cysteines, OvAstC) and tested recombinant OvAstCR1 or OvAstCR2 activation.

The synthetic OvAstC peptides elicited activation of OvAstCR1 and OvAstCR2 in a dose-dependent manner (Figure 6A). A 10-fold shift in the EC_50_ value was observed between the two forms of the mature peptide, with the cyclised version showing higher affinity for both receptors (e.g., AstCR1: OvAstC EC_50_ = 2.8 x 10^-8^ M, 95%CI: 2.0x10^-8^ – 4.1 x 10^-8^ M vs OvAstC_nc EC_50_ = 1.6 x 10^-7^ M, 95%CI: 8.4 x 10^-8^ – 1.3 x 10^-7^ M).

**Figure 6.**
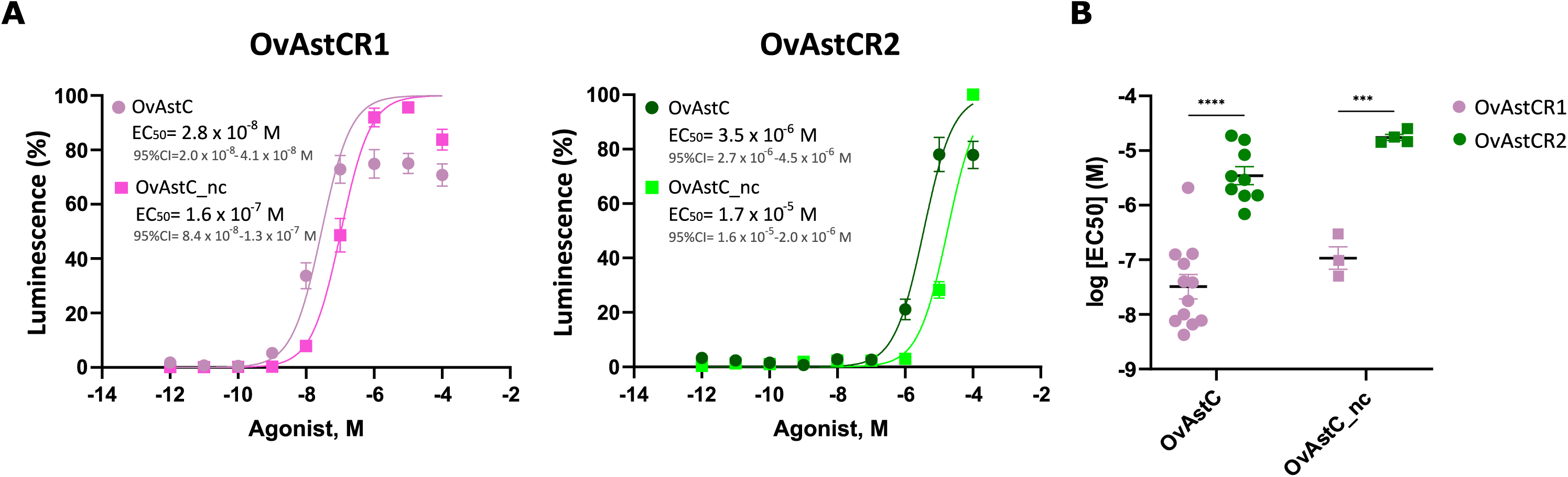
OvAstC activates both OvAstCRs in a dose-dependent manner. ***A*** Dose response curves of OvAstCRs in the presence of the mature cyclic (AVITA**C**YFQAVS**C**Y) and non-cyclic (AVITACYFQAVSCY) AstC peptide expressed as normalised response ± s.e.m. The EC50 value and related 95%CI for each receptor-peptide pair is shown. OvAstC: cyclic form of allatostatin C mature peptide; OvAstC_nc: non-cyclic form of allatostatin C mature peptide. Each indicated peptide-receptor pair was tested in triplicates in each plate and the experiment was repeated at least three times. **B** Plot showing the average EC50 values ± s.e.m of each receptor with the different forms of AstC peptide. There is a 100-fold difference in the EC50 between OvAstCR1 and OvAstCR2 that is visible with both the cyclic and non-cyclic form of the peptide. Comparison between the EC50 values of the different AstCRs with the non-cyclic and cyclic form of the peptide was performed with a two-way ANOVA and Tukey’s multiple comparisons test as a post-hoc analysis. The level of significance was set at p<0.05. ***=p<0.001; ****=p<0.0001.

When comparing the receptors’ affinity for the peptide, a 100-fold difference in the EC_50_ value between OvAstCR1 and OvAstCR2 was observed for both peptides (Figure 6B) with OvAstCR1 showing higher affinity when compared to the OvAstCR2 (e.g., OvAstC: AstCR1 EC_50_ = 2.8 x 10^-8^ M, 95%CI: 2.0 x 10^-8^ – 4.1 x 10^-8^ M vs AstCR2 EC_50_ = 3.5 x 10^-6^ M, 95%CI: 2.7 x 10^-6^ – 4.5 x 10^-6^ M, ****p<0.0001).

## Discussion

### *O. vulgaris* possess two allatostatin C receptors with a tissue distribution consistent with a broad physiological role

In this study we identified for the first time, a putative allatostatin C signalling system in the cephalopod mollusc *O. vulgaris* using *in silico* and functional approaches.

GPCRs are characterised by a very conserved membrane-spanning structure, and the specific receptor identity can only be resolved by complementing multiple alignments, genome structural analysis and phylogenetic investigation with functional studies that ultimately demonstrate the interaction with the predicted ligands (Yañez Guerra and Zandawala, 2023). Our previous bioinformatic analysis of *O. vulgaris* transcriptome, identified two putative AstCRs (Pieroni *et al.*, 2026). However, while sequence alignment and secondary structure prediction analysis showed unequivocable conserved signatures of class A GPCRs such as DRY and NSxxNPxxY motifs (Figure 1), cluster-based and phylogenetic investigations were carried out to confirm their classification as AstCRs.

Indeed, our phylogenesis showed OvAstCR1 and OvAstCR2 to be the only two *O. vulgaris* representatives identified in the AstCR invertebrate-specific branch (Figure 3). In line with what has been previously shown, AstCRs are the closest protostome orthologues of the vertebrate-specific OPRs and are paralogous receptors of the deuterostome-specific somatostatin (Mirabeau and Joly, 2013; Elphick, Mirabeau and Larhammar, 2018; Zhang *et al.*, 2022). Interestingly, our phylogenesis suggests that AstCRs have undergone a series of internal duplications leading to at least two types of receptors in different lineages. We found a single receptor in Crustacea, Annelida and Gastropoda, but two receptors in Rotifera, Bivalvia, Cephalopoda and Insecta (Figure 3). In particular, the nematode *C. elegans* showed a remarkable family expansion with 8 different encoded AstCRs, characteristic of a one-to-many orthologues relationship relative to other bilaterians (Kuzniar *et al.*, 2008).

The shared phylogeny of AstCRs with members of SSRs and OPRs makes the case for asking whether the molecular similarity is also shared at the functional level. Despite the conserved class A GPCR features, such as the intracellular elements involved in the recruitment and interaction with the cellular G-protein, the extracellular elements that are involved in the formation of the ligand binding site are extremely different across the three subfamilies and seem to have evolved to provide discrete differences in agonist recognition. This is supported by experiments showing that opioid peptides and somatostatins are unable to activate *D. melanogaster* AstCRs and that replacing the fruit fly ligand binding site with residues from the rat SSR2 impaired the activation of DmelAstCR by the endogenous peptide (Kreienkamp *et al.*, 2002).

Although the specificity of the ligand binding site impedes cross reactivity between the different receptors, an overlap in the functions can indeed be found.

The main role identified for AstC is the inhibition of the synthesis of the juvenile hormone in some insects corpora allata (Kramer *et al.*, 1991). To some extent this hormonal regulatory role is retained in vertebrates, where somatostatins have been found to inhibit the release of the growth hormone from the pituitary gland (Brazeau *et al.*, 1973). Furthermore, for both signalling systems and to some extent the opioid system too, the broad tissue distribution has been found to be associated with promiscuous functions in the organism physiology. AstCRs have been found across the digestive system and the musculature of intestine and buccal mass in invertebrates, supporting the identified function in feeding and digestion, with a role both at the motility (with a myo-modulatory role) and at the secretory level (Ma *et al.*, 2009; Stemmler *et al.*, 2010; Alzugaray, Hernández-Martínez and Ronderos, 2016; Kubrak *et al.*, 2022; Zhang *et al.*, 2022). Similarly, SSRs have been found to be implicated in gut motility, pancreatic regulation and digestive secretions (Olias *et al.*, 2004; Tostivint *et al.*, 2014) and OPRs have been reported to be involved in the modulation of fluid and electrolytes transport in the gastrointestinal tract (Sternini *et al.*, 2004). The expression of both OvAstCRs in tissues related to the digestive tract such as stomach, intestine, gastric ganglion and secretory glands like the digestive gland, supports this potential role is conserved in cephalopods as well (Figure 5D).

An immune-related role has been proven for the AstC system in bivalves and suggested by the broader distribution found in the haemolymph of different invertebrate species (Skiebe, 1999; Li *et al.*, 2021). A similar role has been reported for somatostatin and opioids as well (Krantic, 2000; Ferone *et al.*, 2004; Liang *et al.*, 2016). We have been able to amplify the genes of interest from immune-related tissues of octopus, including the haemocytes and the white bodies where they are produced, or the posterior salivary gland where substances (e.g., venom, enzymes or other neuropeptides) used as a defence or protection are secreted from (Gestal and Castellanos-Martínez, 2015).

Perhaps the most widely recognised function of the vertebrate opioid system is its central role in analgesia and in the modulation of nociceptive processing. In this context the potential role of this peptidergic signalling in antinociceptive response in cephalopods becomes particularly relevant, as similar function are observed across AstCRs, OPRs and SSRs (Carlton *et al.*, 2001; Pintér, Helyes and Szolcsányi, 2006; Yaksh and Wallace, 2011). Indeed, the lack of convincing opioid peptides and receptors from invertebrate genomes and tissues has prompted the view that these animals do not possess the capacity to experience pain as defined by the Bateson’s criteria (1991). However, recent evidence has demonstrated that AstC is involved in the suppression of aversive thermal responses in *D. melanogaster*, thus suggesting the presence of a more ancestral analgesic system predating the emergence of the actual vertebrate-exclusive one (Bachtel *et al.*, 2018; Liu *et al.*, 2023). We have detected the presence of *OvAstCR1*, *OvAstCR2* and *OvAstC* in the central brain regions (SEM, SUB, OL) and in peripheral tissues usually associated with sensory responses such as arms and suckers. In cephalopods, the central brain mass comprises interconnected lobes responsible for the integration of sensory information, motor coordination and higher-order processing, including learning and behavioural modulation (Young, 1971). The detection of *OvAstCR1*, *OvAstCR2* and *OvAstC* within these regions therefore suggests that AstC signalling may act centrally to modulate neural circuit activity, rather than being restricted to peripheral effector tissues, suggesting a top-down action through the stimulation of potential descending pathways that is worth exploring in the context of nociceptive stimulation.

Altogether, our gross analysis of the tissue distribution supports a conserved pleiotropic, physiological role of the *O. vulgaris* allatostatin C system, but more detailed studies on the actual localisation of the receptors within the cells, combined with functional and behavioural investigations in the whole animal, are needed to confirm this (Figure 5D).

From a genomic point of view, the *OvAstCRs* were found to be intronless genes, similarly to most of the human SSRs (Tostivint *et al.*, 2014). The genome of *O. vulgaris* reports 87% of transcripts are multi-exonic (Destanović *et al.*, 2023); this is in line with a large occurrence of introns reported in invertebrate genomes, including GPCR A receptors (Bryson-Richardson *et al.*, 2004). When compared to other species in which a genomic analysis has been performed, *OvAstCRs* are similar to the mosquito *Aedes aegypti* AstCRs and other insects from coleoptera, hymenoptera and lepidoptera which are intronless (Mayoral *et al.*, 2010). However this is different from *Drosophila* AstCRs which have two introns each (Kreienkamp *et al.*, 2002).

The evolutionary significance of such occurrence would be interesting to investigate in association with a proposed scenario of local gene duplication and subsequent “dispersion” into the genome. This was also suggested in a previous large scale investigation of cephalopod GPCRs evolution where the AstCRs are found to belong to the unexpanded group of genes which therefore derives from conserved GPCR families (Ritschard *et al.*, 2019).

### Cephalopod AstC mature peptide shows a strong conservation across species and strong similarity across phyla

Even though we found two AstCRs in *O. vulgaris* genome, our *in silico* analysis led to the detection of a single peptide precursor. Apart from two peptides previously characterised in insects (Kramer *et al.*, 1991; Kreienkamp *et al.*, 2002), our results are in line with findings from other molluscan species with two receptors, where a single peptide was identified as well (Li *et al.*, 2021).

The only study in cephalopods that surveyed the neuropeptidome of *S. officinalis* did not report the presence of AstC (Zatylny-Gaudin *et al.*, 2016). However, this could be ascribable to challenges in interpreting cyclised peptides with mass spectrometry rather than the actual absence of the peptide (Chaturvedi, Bawake and Sharma, 2024). In support of this, an *in silico* analysis using our identified OvAstC precursor sequence revealed the presence of a single predicted peptide in the transcript assembly of other cephalopods including *Sepia* (Figure 4B). Interestingly, in all cases analysed, despite slight changes in the signal peptide region, the mature peptide shared a 100% identity across all the cephalopod species (Figure 4B). When compared to other protostome peptides, a strong conservation of the cysteines that are notably involved in the formation of a disulfide bridge and of the surrounding residues could be found, except for the third residue following the first cysteine, which was changed from a highly conserved N into a Q in Cephalopoda (Figure 4B). It would be interesting to experimentally investigate whether this might represent a key cephalopod-specific functional residue in the activation of/interaction with the receptors.

The *OvAstC* precursor sequence was found widely expressed across the tissues analysed, supporting the potential promiscuous role, for instance as a neuromodulator in the sensory tissues and brain, or with hormonal-like function as suggested by the presence in the haemocytes or in the salivary glands (Figure 5E).

### Distinct activation profiles of *O. vulgaris* allatostatin-C receptors revealed by Ca^++^-based deorphanisation

The use of a Ca^++^-sensitive aequorin-based assay demonstrated that the predicted mature form of OvAstC could activate the two OvAstCRs in a dose-dependent manner, leading to the identification of a putative AstC system in *O. vulgaris* (Figure 6A). An interesting result was the finding of a very different affinity for the endogenous peptide between the two receptors. Several reasons could explain this 100-fold difference in their EC_50_ between OvAstCR1 and OvAstCR2 (Figure 6B).

The high conservation of the key intracellular amino acids in class A GPCRs, suggests this difference is unlikely to be found in the G-protein binding site, but rather at the level of the ligand binding region. To this regard, in most of the rhodopsin GPCRs, including SSRs and AstCRs, the ligand binding site is formed by the interaction between the N-terminal region, the ECL2 and ECL3 (Peeters *et al.*, 2011). A recent study carried out in the Indian stick insect *Carausius morosus,* showed that the N-terminal is rather variable and seems to be involved in maintaining the conformational structure of the binding site, while the ECL3 is actually mediating the key peptide interaction (Işbilir *et al.*, 2020). In insects, a conserved IxTPP motif has been found to be conserved, and mutating this motif leads to a strong reduction in potency and efficacy of the peptide action (Duan Sahbaz *et al.*, 2017; Işbilir *et al.*, 2020).

Interestingly, this motif is not particularly recurrent in other phyla such as annelids, gastropods and bivalves, but OvAstCR1, which displayed the highest affinity for the peptide (Figure 6A), does show a similar VxxTPP motif (Figure 1). As this pattern is missing in OvAstCR2, it would be interesting to verify whether in cephalopods these residues might play a significant role, similarly to insects, in the ligand-mediated activation of the receptor.

Another hypothesis is that OvAstCR2 is only able to work efficiently if heterodimerising with OvAstCR1. The overlapping expression of both receptors in the same tissues, although not indicative of the actual cellular distribution, might be supportive of the potential functional association. Fluorescence Resonance Energy Transfer (FRET) studies have reported that a series of class A GPCRs in *D. melanogaster,* including DmelAstCR2, are subjected to the formation of homodimers (Rizzo and Johnson, 2020). Heterodimers within the same family (i.e., with DmelAstCR1) were not tested but interaction with different receptor families did not show any significant FRET associated luminescence (Rizzo and Johnson, 2020).

Data on the vertebrate counterparts, show that SSRs often associate to form heteroligomers with other family members or even other GPCRs such as OPRs and dopamine receptors, with resulting significant changes in their pharmacological properties (Olias *et al.*, 2004).

An alternatively explanation for the receptors’ differential response to the putative endogenous peptide might be a direct consequence of the action of the peptide. As proposed above and as also suggested by other works, OvAstC might have a dual action depending on the tissue: it may act as a neuromodulator, therefore displaying a neurotransmitter-like release, or as a secreted compound, thus showing a more hormone-like release. As a consequence, this might require the receptors to exert a different sensitivity to different secreted concentrations, or to have a different cellular distribution (Mayoral *et al.*, 2010; Muscato *et al.*, 2021). Spatiotemporal activity of the peptide relative to the developmental stage of the organism could also be interesting to investigate in this context (Kreienkamp *et al.*, 2002).

Finally, looking at the non-cyclic vs cyclic form of the peptide, there is no selective activation of the receptors that might suggest a specialisation of both receptors and peptide, and despite the possibility of both forms co-existing in the organism, it is most likely that the cyclised peptide make up the prevalent bioactive physiological form.

### Concluding remarks and future directions

We here presented the discovery of a putative endogenous allatostatin C signalling system in *O. vulgaris.* The detection of a functional neuropeptide system known to be implicated in many biologically relevant functions, constitutes an important contribution to the understanding of the physiology of this class of molluscs in relation to their evolutionary context.

Of particular interest is the potential role this system might have in the organisation of octopus antinociceptive signalling, a pathway that has been for long neglected in invertebrates and only predicted in cephalopods, ultimately leading, following the precautionary principle, to their inclusion in UK and EU legislations that regulate their welfare when used in research (Bradshaw, 1999; European Parliament and Union, 2010; UK Statutory Instruments 2012 No. 3039, 2012). The evolutionary relationship with the vertebrate-exclusive OPRs and the recent experimental evidence of the AstC role in suppressing noxious responses in insects (Bachtel *et al.*, 2018; Liu *et al.*, 2023), make this family of receptors a good candidate for further investigation in this behavioural framework, thus contributing to our understanding of the potential molecular contributors of complex nociception in these species.

## Materials and Methods

### In silico analysis

#### *O. vulgaris* allatostatin C precursors search and manual curation

Allatostatin C peptide precursor sequences from known invertebrate species were retrieved from literature and complemented with those found on Uniprot within the Protostomia (id: 33317). Each query was manually filtered for non-relevant incomplete sequences and subsequently blasted against *O. vulgaris* genome (Destanović *et al.*, 2023) using 10e^-2^ as an e-value cut-off, due to the challenge posed by short peptide precursors sequences to be identified (Baggerman *et al.*, 2005). Sequence identity and similarity were analysed using EMBOSS Water with a BLOSUM62 matrix (Madeira *et al.*, 2024). The list of sequences used is available in Supplementary File S1.

The mature form of the peptide was predicted using SignalP - 6.0 to detect the signal peptide (Nielsen, 2025) and Neuropred to detect the cleavage site in the precursor (Southey *et al.*, 2006). We additionally checked for the presence of two cysteines that are usually involved in the formation of a disulfide bond in this class of peptides (Veenstra, 2010).

#### Genes exon-intron boundaries detection

Receptors and peptide precursor transcript sequences were aligned against the *O. vulgaris* chromosome scaffolds (Assembly: GCA_951406725.2) to find their genomic location using NCBI-Magic-BLAST tool (Boratyn *et al.*, 2019). The results were sorted using samtool (Danecek *et al.*, 2021) and the data was processed with samtool idxstats and visualised using Tablet software (The James Hutton Institute, v1.21.02.08, Milne *et al.*, 2013).

#### Cluster analysis and phylogenetic investigation of OvAstCRs

Reference protein sequences for allatostatin C, opioid and somatostatin receptors were selected from *Drosophila melanogaster*, *Aplysia californica* and *Homo sapiens* based on de-orphanised or curated sequences. Reference sequences for the outgroups galanin, Kiss1 and allatostatin A receptors were also included based on previous works showing their affinity to the target receptors that are the focus of this work (Mirabeau and Joly, 2013; Bauknecht and Jékely, 2015). The proteomes of 21 representative species from different phyla were selected based on the best Benchmarking Universal Single-Copy Orthologs (BUSCO) value (Supplementary Table S1) and used as a template for blasting the reference sequences (e-value 1e^-10^ and 40 maximum target sequences). The results from each species were sorted and processed through CD-hit with a similarity threshold of 0.95 to remove incomplete sequences, shorter isoforms or redundant sequences (Li and Godzik, 2006; Fu *et al.*, 2012). The results containing unique representative sequences from each species (1084, Supplementary File S2) were then subjected to a cluster analysis using the CLuster ANalysis of Sequences (CLANS) toolkit, setting a BLAST HSPs e-value of 1e^-12^ and BLOSUM62 scoring matrix (Zimmermann *et al.*, 2018) with an initial p-value threshold of 1e^-60^, to visually reveal the relationships between the sequences. However, as we were missing already annotated sequences (i.e., *C. elegans* AstCRs; Beets *et al.*, 2023), we reduced the threshold to 1e^-40^. All the sequences of the cluster which included the *O. vulgaris* allatostatin C receptors (190) were selected using the convex clustering option (cutoff std dev: 0.6; min sequences per cluster: 20). Additional 7 sequences from *C. elegans* were included as they showed a direct or transitive connection with the AstCRs subgroup. The final 197 sequences were used to generate a phylogenetic tree (Supplementary File S3).

The sequences were aligned with MAFFT v7.526 using the E-INS-i method (Katoh and Standley, 2013) and then trimmed with TrimAl (Supplementary File S4) using default parameters and gappy-out mode (Capella-Gutiérrez, Silla-Martínez and Gabaldón, 2009). The best-fit model of evolution was selected with ModelFinder (LG+F+R7 chosen according to the Bayesian Information Criterion) in IQ-TREE3 v3.0.1 (Wong *et al.*, 2025). Tree branches were obtained using aLRT-SH-with 1,000 replicates and ultrafast bootstrap method. The final phylogenetic relationship was visualised using FigTree v1.4.4 (http://tree.bio.ed.ac.uk/software/figtree/). The tree was rooted using the outgroups galanin, Allatostatin A and Kiss1 receptors.

### Animal samples

#### *O. vulgaris* sample collection

Three young adults of *O. vulgaris* were caught by local artisanal fishermen in the Bay of Naples, Italy following the local Animal Welfare Body authorisation (Ethical Clearance: ecACR-2302ts36). Animals were weighed, sexed and humanely killed in compliance with the Directive 2010/63/EU (Annex IV) and following best practice protocols described in Andrews et al. (2013), Fiorito *et al.* (2015), Butler-Struben *et al.* (2018). Furthermore, animal-derived materials were utilised in a responsible and ethical way, according to the principles of Replacement, Reduction, and Refinement (3Rs).

The following tissues were collected by a FELASA certified (function D) competent person: supra-oesophageal mass (SEM), sub-oesophageal mass (SUB), optic lobe (OL), stellate ganglion (StG), gastric ganglion (GG), arm (at 50% of its length), tip of the arm (TIP), sucker (Su), digestive gland (DG), anterior salivary gland (ASG), posterior salivary gland (PSG), white bodies (WB), haemocytes (Hc), stomach (Stom), Intestine (Int). Tissues were frozen in 50 μL RNA later and shipped in dry ice. The tissues were processed using TRIzol-RNA extraction and cDNA synthesis (Supplementary Table S2).

### Molecular biology

#### RNA extraction and cDNA synthesis from *O. vulgaris* tissues

Tissues were resected into smaller samples (4-40 mg) and placed in 2mL Eppendorf tubes containing 500 μL TRIzol (Cat No. 15596026, Invitrogen™). Each sample was frozen in liquid nitrogen for a few seconds prior to mechanical homogenisation using the Handheld Homogeniser SHM1 (Cole-Palmer). After a 5 min incubation at room temperature, 200 µl chloroform (Cat No. 200-663-8, Sigma-Aldrich) was added, mixed, and incubated on ice for 15 min. Samples were centrifuged (12,000 x g, 4°C) and the supernatant was transferred to a silica-cartridge for the total RNA purification (PureLink® RNA Mini Kit; Cat No 12183018A, Invitrogen^TM^). The purified samples were processed with DNase I, Amplification Grade (Cat No. 18068015, Invitrogen^TM^) following the manufacturer’s instructions. Quality (260/280 ratio close to 2.0) and quantity of extracted RNA was assessed through UV-visible absorption measurements (NanoDrop™ 2000/2000c Spectrophotometers, Thermo Scientific™). RNA samples were used for reverse transcription with SuperScript™ III Reverse Transcriptase (Cat. No. 18080093, Invitrogen^TM^) in which 1 μg of RNA was utilised to produce cDNA. Octopus RNA, as for all the molluscs, is characterised by the so-called ‘hidden break’ in 28S rRNA causing the 18S and 28S rRNA fragments to co-migrate electrophoretically as a single band, thus challenging resolution of the RNA quality through a gel (Natsidis *et al.*, 2019; Adema, 2021). Given this, only cDNA derived from extracted tissues which successfully amplified our control gene *cullin 1* (Imperadore *et al.*, 2023) was selected to be further used in endpoint PCR to evaluate expression of the indicated genes of interest (Supplementary Table S2).

#### Primer design and end point PCR

Primer sequences of *OvAstCR1*, *OvAstCR2* and *OvAstC* were designed using ApE-A plasmid Editor© v3.1.3 software and purchased from Eurofins Genomics (Supplementary Table S2). Once diluted to a working concentration of 25 pmol/µL, the primers were used to amplify the target genes in the tissues described above. PCR amplifications were performed using Phusion™ High-Fidelity DNA Polymerase (Cat No. 10024537, Thermo Scientific™) following manufacturer’s instructions and the setup was adjusted according to the amplicon length and melting temperature of each primer pair. A volume of 1 μL of cDNA synthesis reaction was used for each amplification.

#### Plasmid and peptide synthesis

The cyclic and non-cyclic forms of *O. vulgaris* mature allatostatin C peptide (AVITACYFQAVSCY) were synthesised using the custom peptide synthesis service (PeptideSynthetics, Peptide Protein Research Ltd., UK); The predicted *O. vulgaris* Allatostatin C receptors *OvAstCR1* and *OvAstCR2* were codon optimised for *H. sapiens* expression (Supplementary File S5) and synthesised into the pcDNA3.1+ vector using Genscript gene synthesis service (Genscript Biotech, UK).

#### *O. vulgaris* AstCRs de-orphanisation

The cell line utilised in this study was the HEK293G5A, stably expressing the chimeric GFP-Apoaequorin protein G5A (Cat Nr. cAP-0200GFP-AEQ-Cyto, Angio-Proteomie). Cells were grown in a T75 flask at 37 °C in 5% CO_2_ atmosphere in DMEM (containing 4.5 g/L glucose, L-Glutamine, sodium pyruvate, Cat. No. 10566016 ThermoFisher™) supplemented with 10% FBS (heat inactivated, Cat. No. 10082147, ThermoFisher™) and broad-spectrum antibiotic-antimycotic (Cat No. 15240062, Gibco^TM^). When the culture reached confluency, cells were detached from the bottom of the flask by incubation with 5 mL of TrypLE™ (Cat No. 12605010, Gibco™) for 5 min. The suspension was then centrifuged (1,500 rpm for 5 min), the pellet was resuspended in DMEM (1:5 volumes) prior transfer into Corning™ 96-Well, Cell Culture-Treated, Flat-Bottom Microplates (Cat No. 10517742, ThermoFisher™) and grown for 2 days at 37 °C in 5% CO_2_ atmosphere. For the transfection, for each well, 10 µl OptiMEM (without FBS, Cat No. 11058021, Gibco™), 60 ng of GPCR-containing plasmid, 60 ng of Gqi5/9 plasmid, and 0.30 µg per 100 ng DNA of 25 kDa linear polyethylenimine (PEI; Cat. number: 23966 PolySciences, Inc.) were mixed and incubated for 20 min at room temperature prior adding the mix to 90 µl of Advanced DMEM (Cat No. 12491015, Gibco™) and finally replacing the old medium in the 96-well plate. The transfected culture was incubated at 37 °C and 5% CO_2_ for two days. On the experimental day, the medium was removed and substituted with HBSS media (Cat. No. 20011, AAT-Bioquest) supplemented with 4 µM coelenterazine-H to reconstitute apoaequorin into functional Ca^++^-sensitive aequorin (Cat. No. S2001, Promega). The plate was covered with aluminium foil to protect it from light exposure and incubated at 37 °C in 5% CO_2_ for 2 h. Readings were performed with a FlexStation 3 Multi-mode Microplate reader (Molecular Devices, LLC, USA) using the Flex option, for a period of 75 s per well and ligand injection after 15-18 s.

Each indicated peptide-receptor pair was tested in triplicates in each plate and the experiment was repeated at least three times. Cells transfected with the empty vector alone (pcDNA 3.1) were tested to check for endogenous activation. Cells viability was assessed by administering ATP and checking for a strong comparable activation across the wells.

#### Normalisation and statistical analysis

Data were normalised to the maximum response obtained in each experiment (100% activation) and to the luminescent background recorded in the media alone (0% activation). Dose–response curves were fitted with a four-parameter curve and EC_50_ values were calculated from dose–response curves based on at least three measurements from three independent transfections. Comparison between the EC_50_ values of the different AstCRs with the non-cyclic and cyclic forms of the peptide was performed with a two-way ANOVA and Tukey’s multiple comparisons test as a post-hoc analysis. Data were analysed using GraphPad Prism version 10 for Windows (GraphPad Software, Boston, Massachusetts, USA).

## Data Availability statement

Supplementary Tables S1-S3 and Supplementary Files S1-S5 are electronic files accessible from the following link: **<u>10.5281/zenodo.18067437</u>.** All data generated or analysed during this study are included in the Supplementary Information. Any additional information required is available from the corresponding author upon request.

## Acknowledgments

This work was supported by the HSA-Ceph 1/2019 grant to the Association for Cephalopod Research ‘CephRes’ ETS, Napoli, Italy, and The Gerald Kerkut Charitable Trust, UK.

